# Top-down processing alone activates the early somatosensory nuclei

**DOI:** 10.64898/2026.04.21.719886

**Authors:** Paige Howell, Lynn Farner, Finn Rabe, Tim Emmenegger, Maryam Seif, Patrick Freund, Nicole Wenderoth, Sanne Kikkert

## Abstract

The early somatosensory brainstem and thalamic nuclei are classically viewed as relays of peripheral input, yet research in animal models indicates they also receive top-down cortical signals. Whether such processing exists in humans remains unknown. Using cervical spinal cord injury (SCI), in which peripheral somatosensory input is reduced or absent while cortical processing is preserved, we tested whether top-down processing can elicit activation across the somatosensory nuclei.

We combined 3 Tesla functional and quantitative MRI data to assess activity and structural properties along the somatosensory hand pathway in a cross-sectional study of 16 individuals with chronic cervical SCI (mean age ± s.e.m.=52.4 ± 3.5 years) and 20 age-, sex-, and handedness-matched able-bodied control subjects (mean age=50.8 ± 3.5 years). Participants were visually cued to make overt or, in cases of hand paralysis, attempted right- and left-hand movements. Activation was quantified across the cuneate nucleus, ventroposterior lateral thalamus, and primary somatosensory hand cortex, while structural properties were assessed using quantitative MRI measures sensitive to myelin and tissue integrity, including magnetisation transfer saturation (MTsat) and effective transverse relaxation rate (R2*), alongside morphometric measures.

Despite reduced or absent peripheral input, SCI participants exhibited robust and lateralised activation across all levels of the somatosensory pathway. This pattern persisted even in a participant with complete hand paralysis who lacked bottom-up afferent input during the fMRI task, indicating that top-down processing alone is sufficient to drive activity in early somatosensory relays. We simultaneously observed structural degeneration in the cuneate nucleus of SCI participants, marked by reduced volume and myelin-sensitive metrics (MTsat and R2*), consistent with secondary degeneration. The extent of atrophy was related to time since injury and reduced sensorimotor hand function, but showed no significant relationship with functional activation, suggesting that preserved corticocuneate signalling is not dependent on the degree of structural degeneration.

This provides the first evidence that the cuneate nuclei in humans are subject to both bottom-up and top-down somatosensory processing. Although these nuclei are vulnerable to structural atrophy following dorsal column injury, our results suggest that top-down processing remains intact decades after SCI. This may have implications for the development of rehabilitation treatments targeting preserved somatosensory processing after injury.

## 1. Introduction

The ability to sense with our hands is crucial for interacting with our environment. This relies on a precisely organised somatosensory system that conveys tactile and proprioceptive input from the periphery to the brain. These afferent signals first synapse in the ipsilateral cuneate nucleus of the brainstem, before projecting to the contralateral ventroposterior lateral (VPL) thalamus, and finally to the hand area of the primary somatosensory cortex (S1; Figure 1).^1^

**Figure 1.**
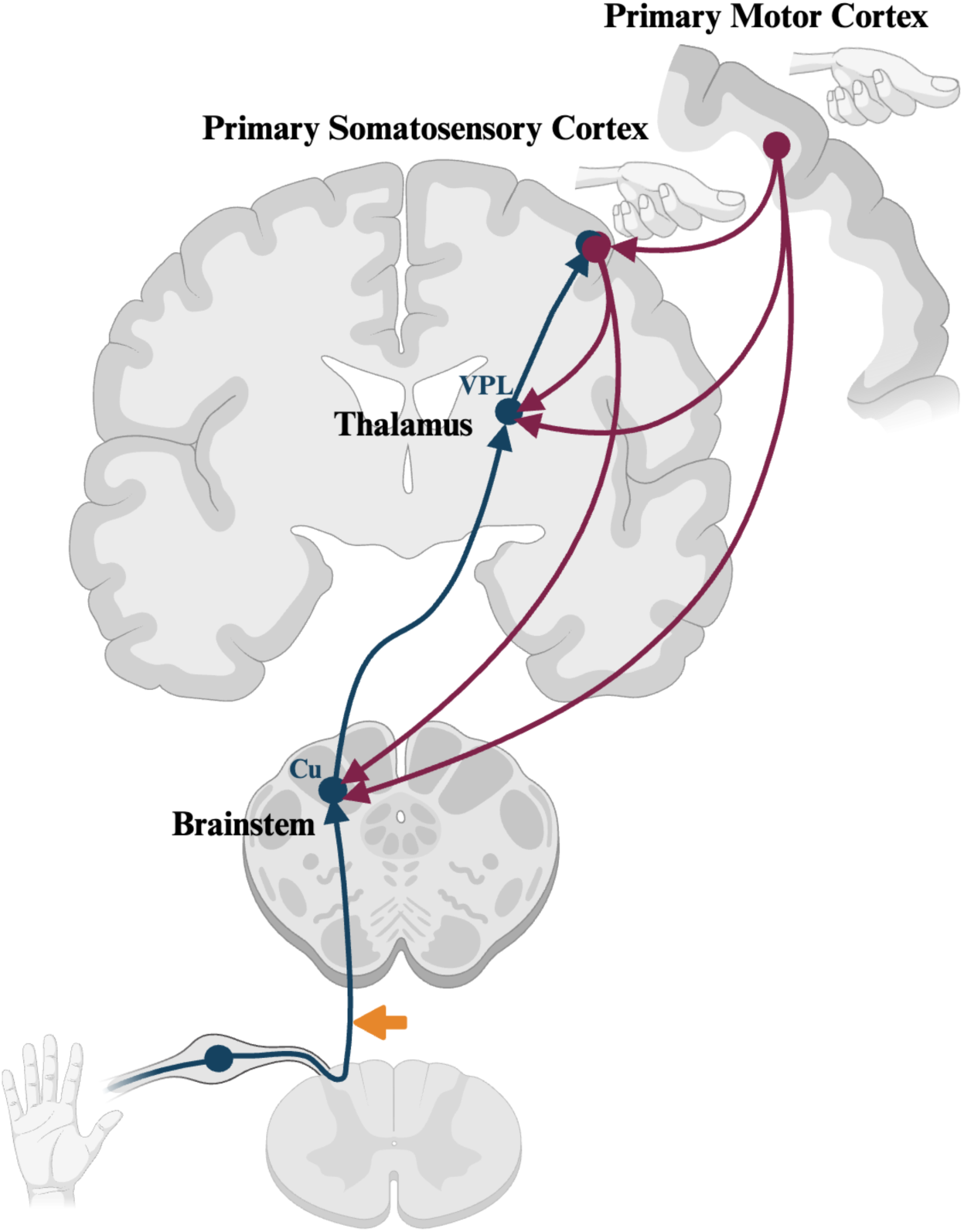
Canonical hand somatosensory pathway and top-down projections from the sensorimotor cortex. Schematic illustration of the somatosensory processing stream. The bottom-up somatosensory pathway (blue) conveys tactile and proprioceptive input from the hand to the ipsilateral cuneate nucleus (Cu). From there, signals transfer to the contralateral ventroposterior lateral (VPL) thalamic nucleus, and finally the hand area of the contralateral primary somatosensory cortex (S1). Top-down projections from the sensorimotor cortex (red), as described in animal studies, are shown projecting to each level of the somatosensory pathway. Following cervical spinal cord injury, the transmission of bottom-up input from below the lesion level (orange arrow) is partially or completely interrupted, providing a model to explore top-down processing in vivo. Figure created with BioRender.com.

Although the early somatosensory nuclei have classically been viewed as mere relay stations for peripheral inputs, previous research in animal models has shown that they also receive top-down projections from the sensorimotor cortex.^2–5^ During active movement, these corticocuneate pathways are thought to modulate the transmission of sensory signals passing through the cuneate nucleus.^3,6,7^ The cuneate nuclei are therefore often termed the somatosensory gateway to the brain.^8^ These corticocuneate projections remain preserved in non-human primates after spinal cord injury (SCI) and have been suggested to contribute to functional recovery.^2^ Despite their crucial role in sensory processing and potential relevance for rehabilitation, it has been extremely challenging to investigate these nuclei in vivo. As a result, knowledge about corticocuneate projections stems exclusively from research in mice^6^, cats^3,5,9–11^, and non-human primates^2,12–14^. Whether such processing exists in humans, and what happens to these projections after sensory loss, remains unknown.

A fundamental challenge in resolving this question is separating top-down somatosensory processing from peripheral sensory input. During voluntary hand movements, top-down activity in the cuneate nuclei is confounded by concurrent bottom-up input generated by the movement itself ^12^, making it difficult to isolate. Cervical SCI disrupts this coupling, as bottom-up afferent inputs from the hand are reduced or absent, while top-down, corticocuneate, processing may remain intact. SCI, therefore, provides a unique model for probing top-down corticocuneate processing.

We previously showed that individuals with complete cervical SCI, i.e., who lack sensorimotor hand function and exhibit a complete disconnect between the brain and the periphery, can activate somatotopic hand representations in S1 during attempted finger movements.^15^ This demonstrates that intracortical processes alone can drive somatotopic activity in S1. Such preserved representations have been leveraged in brain-computer interface approaches to restore somatosensation after deafferentation.^16–18^ Likewise, if corticocuneate projections exist in humans and persist after cervical SCI, then attempted hand movements should elicit activity in the cuneate nuclei. However, given that chronic loss of bottom-up sensory input is associated with structural degeneration of the cuneate nuclei^19–21^, it is unclear to what extent such top-down modulation can persist following SCI.

In this study, we recruited individuals with cervical SCI, in whom bottom-up somatosensory inputs are partially or completely absent, to test whether top-down cortical processing alone can engage somatosensory nuclei throughout the somatosensory processing stream. Using 3 tesla (3T) functional MRI (fMRI), we measure activity in the cuneate nuclei, VPL, and S1 hand area during overt or attempted hand-movement. To evaluate whether top-down modulation of these nuclei persists in the presence of neurodegeneration, we assess the structural properties of these regions using 3T quantitative MRI (qMRI) with multi-parameter mapping (MPM) of macro- and micro-structural metrics. Finally, we examine clinical and behavioural correlates of preserved structure and function to explore factors that may support residual somatosensation after SCI.

## 2. Results

### Patient impairments

We tested 16 individuals with chronic cervical SCI who varied in injury completeness (AIS A to D), neurological level of injury (C1 to C7), time since injury (6 months to 37 years), and hand sensorimotor impairment (bilateral GRASSP hand scores ranging from 3 to 84 out of a maximum possible score of 148). We also tested 20 healthy control participants matched for age, sex, and handedness.

### Relay nuclei along the somatosensory pathway are active despite reduced or absent afferent drive

We assessed whether overt or attempted hand movements after cervical SCI activate cortical and subcortical nodes along the somatosensory processing stream. As expected, whole-brain control-group activity maps demonstrated that overt hand movements elicited robust activation throughout the somatosensory processing stream, consistent with the canonical spatial organisation of this pathway (Figure 1). We observed significant activation clusters overlapping the brainstem cuneate nuclei, the VPL thalamus, and the S1 hand area (Figure 2A).

**Figure 2.**
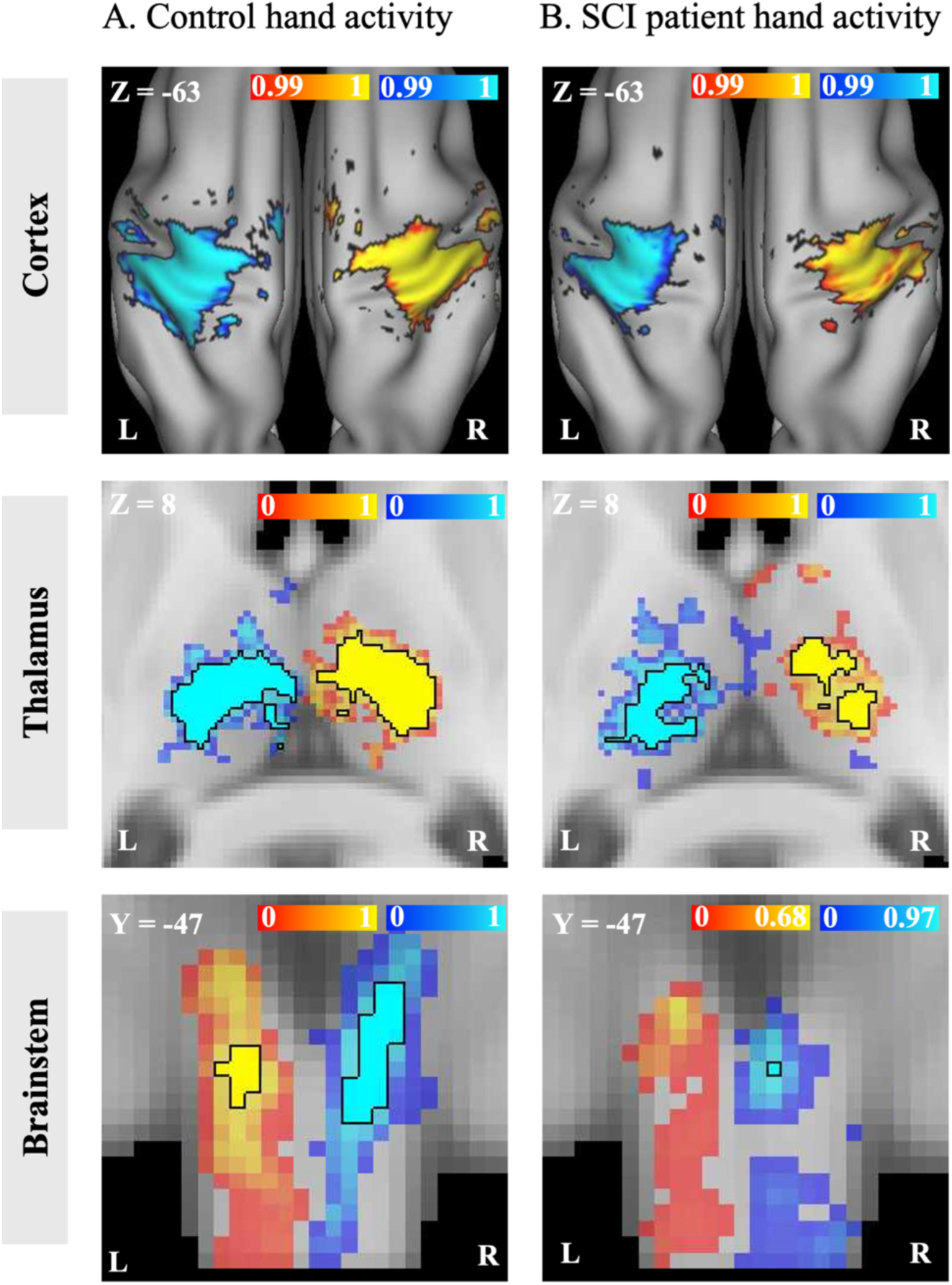
Group-level activation during hand movement follows the canonical somatosensory pathway in controls and cervical SCI. Group-level activity maps for control participants (A) and individuals with cervical spinal cord injury (B) during (attempted) hand movements. Both groups showed activation clusters overlapping with the key relay stations along the somatosensory processing stream (see Figure 1 and Figure 3A), including the brainstem cuneate nuclei, the ventroposterior lateral (VPL) thalamus, and the S1 hand area. Suprathreshold clusters are outlined in black, with subthreshold activity displayed at a reduced opacity. Colour bars represent 1 − pFWE (family-wise error- corrected p-values). Warm colours (red–yellow) represent the contrast non-dominant > dominant hand (left panel) and more impaired > less impaired hand (right panel), whereas cool colours (light blue–blue) represent the inverse contrasts (dominant > non-dominant hand; less impaired > more impaired hand).

To quantify activation magnitude and laterality within the hand somatosensory nuclei, we performed ROI analyses at each pathway level using predefined anatomical ROIs (Figure 3A). We averaged activity across both hands to derive ipsilateral and contralateral responses relative to the moving hand. This analysis confirmed strong lateralisation across all regions, with greater activity in the ipsilateral cuneate nucleus (t_(34)_ = 8.94, pFDR < 0.001, BF_10_ = 1.4 × 10^5^), the contralateral VPL (t_(34)_ = -10.50, pFDR < 0.001, BF_10_ = 4.9 × 10^5^) and the contralateral S1 hand area (t_(34)_ = -14.48, pFDR < 0.001, BF_10_ = 3.3 × 10^8^) compared with their opposite-side homologues (Figure 3B).

**Figure 3.**
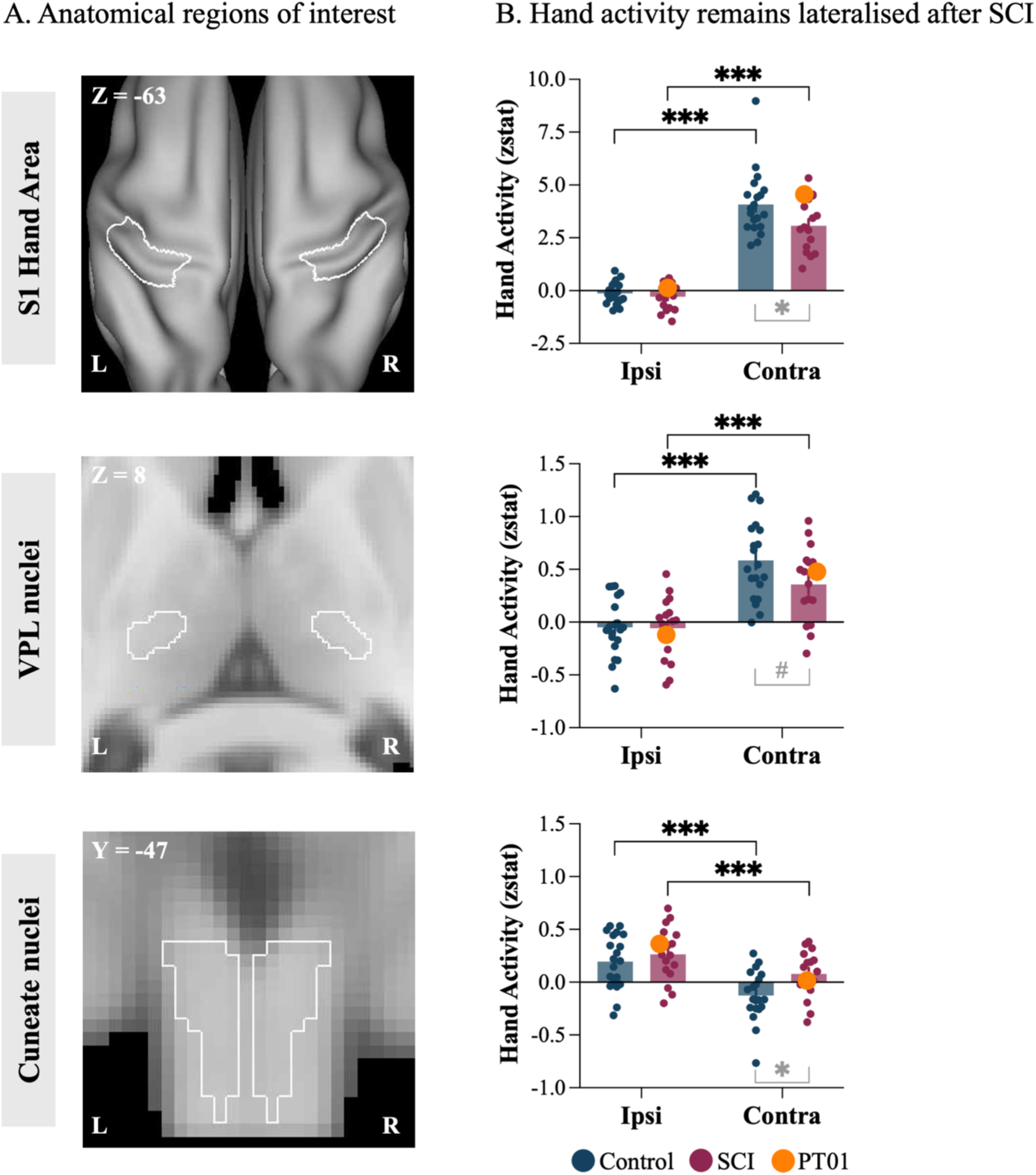
Overt and attempted hand movements elicit lateralised activity across the entire somatosensory processing stream. (A) Anatomical regions of interest (ROIs) along the somatosensory processing stream are outlined in white, showcasing the main relay stations of somatosensory afferent inputs, bilaterally. From top to bottom: the primary somatosensory cortex (S1) hand area, the ventroposterior lateral (VPL) thalamic nuclei, and the cuneate nuclei in the brainstem. (B) ROI analyses show mean activation (z-statistic) during hand movement, for the ROI on the same side as the moving hand (Ipsi) and for the opposite side (Contra). Each plot corresponds to the ROIs shown in the same row of panel. Individual control participants are plotted as blue dots, and individual SCI participants as red dots. The most impaired SCI participant (PT01), who had an anatomical and clinically complete cervical injury, i.e. no spared spinal tissue bridges, and a complete hand paralysis with no detectable EMG during voluntary hand movement (see Supplementary materials), is shown in orange. Across all three ROIs, activity was consistently stronger in the hemisphere corresponding to the moving hand, revealing robust lateralisation of the somatosensory pathway. This lateralised pattern was present not only in controls performing overt movements but also in SCI participants performing purely attempted hand movements (i.e., PT01). p-values are false discovery rate corrected for multiple comparisons within each ROI. *** = p < 0.001, ** = p < 0.01, * = < 0.05.

We next asked whether this canonical pattern was preserved in SCI, despite partial or complete hand paralysis. Participants unable to perform any overt hand movement were instructed to attempt the movement, meaning they generated the intention to move despite no visible muscle activity. In this group, afferent input from the hand during the movement task is reduced or absent, whereas top-down, corticocuneate influences are expected to remain intact. Strikingly, the somatosensory hand pathway remained robustly and selectively engaged in participants with cervical SCI. The overall spatial organisation of the group map activity closely resembled that of the canonical somatosensory pathway and the control group (Figure 2B). Hand activity in cervical SCI participants was strongly lateralised across all levels of the somatosensory processing stream, as demonstrated by a main effect of Side in all three regions (Cuneate: F(1,34) = 88.37, p < 0.001, η²ₚ = 0.72, BF_10_ = 5.2 x 10^7^; VPL: F(1,34) = 134.33, p < 0.001, η²ₚ = 0.80, BF_10_ = 8.5 x 10^10^; S1: F(1,34) = 300.53, p < 0.001, η²ₚ = 0.90, BF_10_ = 4.7 x 10^22^). However, the relative difference between ipsi- and contralateral activity was larger in controls than in SCI participants (Group × Side interactions in each ROI: Cuneate: F(1,34) = 6.35, p = 0.02, η²ₚ = 0.16, BF_10_ = 3.4; VPL: F(1,34) = 5.81, p = 0.02, η²ₚ = 0.15, BF_10_ = 2.8; S1: F(1,34) = 3.88, p = 0.06, η²ₚ = 0.10, BF_10_ = 1.5).

No significant differences in group activity levels were observed in the ipsilateral cuneate nucleus, ipsilateral VPL, or ipsilateral S1 hand area (Cuneate: t_(41.93)_ = -0.83, pFDR = 0.41, BF_10_ = 0.41; VPL: t_(47.21)_ = 0.07, pFDR = 0.94, BF_10_ = 0.32; S1: t_(63.38)_ = 0.42, pFDR = 0.67, BF10 = 0.41; Figure 3B). By contrast, SCI participants showed higher activity in the contralateral cuneate nucleus activity compared with controls (t_(41.93)_ = -2.47, pFDR = 0.02, BF_10_ = 4.0), but lower activity in the contralateral VPL (t_(47.21)_ = 2.05, pFDR = 0.06, BF_10_ =1.2) and contralateral S1 hand area (t_(63.38)_ = 2.80, pFDR = 0.01, BF_10_ = 1.8). Notably, activity in the cuneate was significantly greater than zero in both SCI participants (t_(15)_ = 4.03, pFDR < 0.01, BF_10_ = 35.43) and controls (t_(19)_ = 3.38, pFDR < 0.01, BF_10_ = 13.64).

Because the cervical SCI cohort was heterogeneous, some participants retained limited hand motor function and residual afferent input during the task. We therefore asked whether the same pattern of results would be observed in an individual with an anatomically and clinically complete cervical SCI, with no spared spinal tissue bridges at the lesion level, and a complete hand paralysis, i.e., no detectable hand muscle activity during voluntary hand movements (see Supplementary Materials for EMG testing results). This participant could only perform attempted, rather than overt, hand movements. Therefore, if activity in the cuneate and VPL nuclei would depend on peripheral bottom-up input, the absence of movement, and associated absent afferent feedback, should eliminate this response. Instead, we found preserved and lateralised activity in this participant (PT01; Figure 3B, orange marker) with activation levels that were comparable the control group across all ROIs (Crawford t-test: Cuneate: t_(19)_ = 0.64, p = 0.53; VPL: t_(19)_ = -0.28, p = 0.78; S1: t_(19)_ = 0.30, p = 0.77). Together, these results indicate that top-down cortical signals alone are sufficient to drive activity in the brainstem and thalamic somatosensory nuclei in humans after cervical SCI.

### Structural changes in the brainstem after SCI

A loss of afferent input after cervical SCI is known to trigger structural degeneration within subcortical somatosensory nuclei in both humans and non-human primates.^19,21,22^ To assess structural changes in our SCI cohort and their potential relationship to the observed functional preservation, we next examined structural integrity within the somatosensory regions.

We first compared macrostructural differences between cervical SCI participants and controls using voxel-wise comparisons of white- and grey-matter probability maps to identify spatially localised effects across the whole brain. The white-matter analysis revealed a focal reduction in tissue volume in the dorsal medulla, spatially overlapping with our cuneate nuclei ROI (Figure 4A). We detected no significant clusters in the VPL or S1. This pattern suggests demyelination in the dorsal column nuclei, consistent with anterograde fibre degeneration known to follow cervical SCI.^21,23^ A grey-matter analysis revealed no significant group differences across the brain.

**Figure 4.**
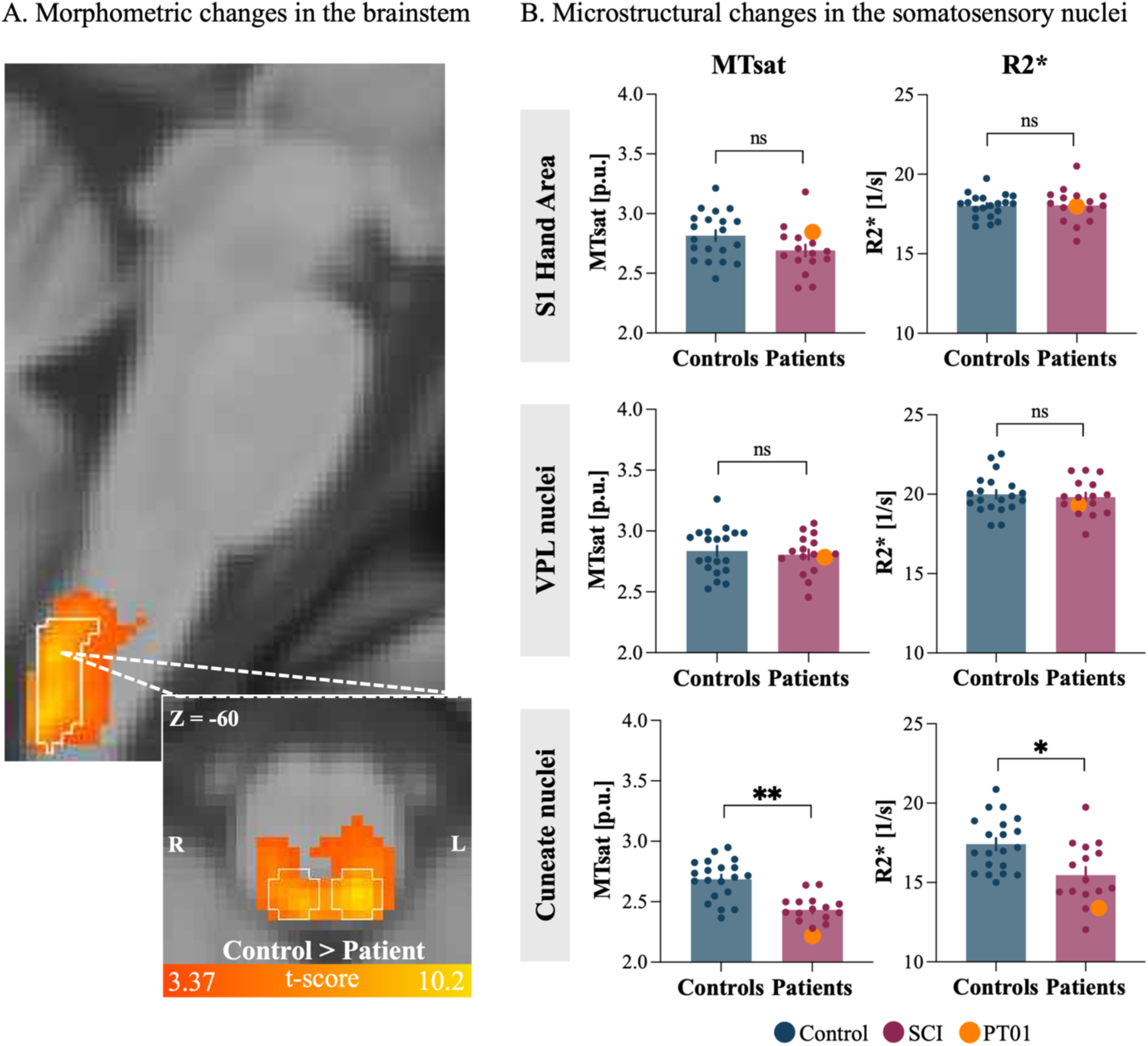
Structural degeneration within the cuneate nuclei of the brainstem after cervical spinal cord injury. (A) We found macrostructural volume loss in SCI participants compared with healthy controls (statistical parametric map thresholded at p < 0.001 voxelwise, uncorrected with cluster-level pFWE < 0.05). Volume loss was localised to the dorsal medulla and overlapped with the cuneate nuclei ROI (outlined in white for visualisation only). (B) Region-of-interest (ROI) analyses showing average quantitative MRI (qMRI) values sensitive to myelin (Magnetisation Transfer Saturation; MTsat) and broader tissue microstructure (R2*) within the S1 hand area, VPL, and cuneate nuclei, bilaterally. SCI participants showed significantly reduced MTsat and R2* values in the cuneate nucleus compared with controls, whereas no significant group differences were observed in the VPL or S1. Individual participants are plotted as blue (controls) or red (SCI) dots. As in Figure 2, a complete SCI participant (PT01) is shown in orange for visualisation (ASIA A, injury level C4, GRASSP score = 0; tissue bridges = 0). Error bars denote the standard error of the mean. p.u. = percentage units; 1/s = per second. *** = p < 0.001, ** = p < 0.01, * = < 0.05.

For a region-specific assessment, we next quantified the microstructural tissue properties within our predefined anatomical masks of the cuneate nuclei, VPL, and S1 hand areas (Figure 3A). We extracted quantitative MRI measures sensitive to myelin content (MTsat) and to changes in tissue composition, including iron (R2*; Figure 4B), and averaged values across homologous ROIs. Within the cuneate nucleus, group comparisons revealed reductions in both MTsat (t_(33.48)_ = 5.40, pFDR < 0.001, d = 1.78, BF_10_ = 1.6 × 10³) and R2* (t_(29.75)_ = 3.11, pFDR < 0.01, d = 1.05, BF_10_ = 12) in SCI participants compared with controls. We found no significant group differences in the S1 hand area or the VPL for MTsat (S1: t_(32.35)_ = 1.87, pFDR = 0.14, d = 0.63, BF_10_ = 1.2; TH: t_(33.84)_ = 0.50, pFDR = 0.66, d = 0.17, BF_10_ = 0.36) and R2* (S1: t_(25.52)_ = −0.07, pFDR = 0.94, d = −0.03, BF_10_ = 0.32; TH: t_(33.15)_ = 0.45, pFDR = 0.66, d = 0.15, BF_10_ = 0.35). Together, quantitative MRI revealed both macro- and microstructural degeneration localised within the cuneate nuclei of our SCI cohort.

### Clinical predictors of preserved structure and function

Finally, we explored which factors may predict preserved activation and/or structural integrity along the somatosensory pathway in our SCI cohort. We first examined whether the magnitude of activation (mean zstat) evoked during the hand-movement task was related to the underlying microstructural integrity within each region (MTsat, R2*). Across the cuneate nucleus, VPL thalamus and S1 hand area, there were no significant correlations between MTsat or R2* values and the corresponding activity estimates (|rₛ| ≤ 0.22, all p ≥ 0.42, uncorrected; Supplementary Figure 3), suggesting no significant linear relationship between these structural and functional measures.

We then investigated whether clinical markers reflecting residual afferent pathways (i.e., tissue bridge width), injury chronicity (i.e., time since injury), or overall hand function (i.e., GRASSP scores) accounted for the task-evoked activation. Across all ROIs, none of our clinical measures explained significant variability in task-evoked activity (full model: Cuneate, R² = 0.19, F(3,9) = 0.71, p = 0.57; VPL, R² = 0.04, F(3,9) = 0.13, p = 0.94; S1, R² = 0.06, F(3,9) = 0.20, p = 0.89). Because tissue bridge measurements were available for only a subset of participants (n = 13), we also ran regression models excluding this predictor, which did not alter the conclusions (reduced model: Cuneate, R² = 0.02, F(2,13) = 0.13, p = 0.88; VPL, R² = 0.14, F(2,13) = 1.10, p = 0.37; S1, R² = 0.07, F(2,13) = 0.52, p = 0.61). This supports the interpretation that the subcortical activation after cervical SCI cannot be explained solely by spared bottom-up afferent signals.

We then explored whether the same clinical measures were instead related to the structural integrity measures across regions. Within the cuneate nucleus, the models predicted both MTsat and R2* measures (full models: MTsat, R² = 0.56, F(3,9) = 3.85, p = 0.05; R2*, R² = 0.55, F(3,9) = 3.70, p = 0.05). This pattern was maintained after excluding the preserved tissue bridge measurements (reduced models: MTsat, R² = 0.47, F(2,13) = 5.66, p = 0.02; R2*, R² = 0.33, F(2,13) = 3.17, p = 0.08). MTsat was significantly associated with retained hand function (GRASSP; full model: B = 0.004, p = 0.02; reduced model: B = 0.003, p = 0.02) and time since injury in the reduced model (full model: B = 0.005, p = 0.12; reduced model: B = 0.006, p = 0.01). R2* values showed a similar pattern, with a significant association with time since injury in the reduced model (full model: B = 0.107, p = 0.07; reduced model: B = 0.096, p = 0.03). Full model statistics for each ROI are reported in Supplementary Table 1.

Together, these findings indicate that the selected clinical scores relate to the degree of structural degeneration in the somatosensory nuclei, whereas no significant associations were observed between these measures and task-evoked activation.

## 3. Discussion

We show that overt and attempted hand movements in cervical SCI participants elicit activity in the brainstem and thalamic somatosensory nuclei, even when peripheral afferent input is severely reduced or absent. This provides the first evidence in humans that top-down cortical processing can drive activity in the early somatosensory relay nuclei.

Despite decades of sensorimotor hand impairment and reduced or absent bottom-up peripheral inputs, cervical SCI participants exhibited clear laterality in processing left- and right-hand signals across the somatosensory relay nuclei. This pattern was consistent with that observed in control participants. Although the activation magnitude in the thalamus and S1 was reduced compared with controls, this reduction is not unexpected, as control responses necessarily reflect both intact bottom-up and top-down somatosensory signalling. Crucially, preserved laterality was observed even in a participant with a complete disconnect between the brain and the periphery, in whom attempted hand movements elicited activity comparable to controls performing overt hand movements. Together, these findings support prior evidence of preserved cortical hand representations following cervical SCI^15,16^ and show that this preservation extends to subcortical levels of the somatosensory system. Moreover, these findings align with post-mortem tracing work in non-human primate SCI models, which shows that cortical projections to the cuneate nuclei can be preserved following both complete and incomplete experimentally induced injuries.^2^

Several neural pathways could contribute to the preserved activation observed in our SCI cohort. Retrograde tracer studies in animal models show that the cuneate nucleus receives dense descending innervation from multiple cortical regions, including motor, premotor, parietal, and cingulate regions.^2,13,24–26^ These pathways have been found to support sensory gating and somatotopic tuning of bottom-up input.^3,6,7^ Attempting hand movements may evoke corollary discharge or efference copy signals that resemble the expected sensory consequences of the action^27,28^, thereby generating top-down sensory activity even when no movement occurs. Furthermore, attempted hand movements may impose high attentional demands, eliciting somatotopic activity in S1^29^, which could, in turn, be relayed to subcortical nuclei through top-down pathways. Importantly, while residual afferent input cannot be excluded in most SCI participants^30,31^, it cannot account for the responses in our most impaired participant. Given that they were unable to generate overt hand movements, no task-related reafferent input could be produced, and even partially preserved bottom-up pathways would have carried little or no meaningful sensory information. Thus, the activation observed in PT01 can be attributed to top-down cortical signalling rather than to afferent drive.

Beyond demonstrating top-down cortical modulation, our findings reveal concurrent degeneration of the cuneate nuclei. Due to their proximity to the injured spinal cord, the brainstem somatosensory nuclei are highly vulnerable to secondary neurodegeneration after SCI, driven by ongoing inflammatory processes and anterograde (Wallerian) axonal degeneration.^32,33^ We observed reductions in white matter volume, MTsat, and R2*, consistent with previous investigations in chronic SCI within the upper cervical cord and brainstem.^20,21,34^ This pattern is suggestive of demyelination and altered tissue composition, associated with the chronic loss of bottom-up afferent input.^35–37^

Despite this structural degeneration, top-down processing appears to remain remarkably preserved after cervical SCI. This dissociation between preserved top-down function and structural loss suggests that degeneration of cuneate tissue reflects disruption of the periphery-brain axis (e.g. residual afferent input, tissue bridges), rather than a failure of corticocuneate signalling. Consistent with this interpretation, we found that structural degeneration scaled with retained hand function (i.e., GRASSP) and chronicity (i.e. time since injury), but not with functional activity. While these findings should be interpreted with caution, considering the limited sample size for multiple regression analyses, the observed associations between clinical measures and structural metrics are consistent with prior work. It has been shown that after SCI, remote neurodegeneration in the upper cervical cord and brainstem scales with impairment, including sensorimotor function^20,37,38^ and time since injury^34,39^.

Given the proposed role of top-down projections in sensory gating, preserved corticocuneate signalling following SCI may provide a promising target for rehabilitation strategies aimed at supporting reafferentation. It has been suggested that top-down modulation could support functional recovery by enhancing the transmission of spared spinal inputs to the cuneate nucleus.^2,19^ Consistent with this idea, studies in non-human primate and rodent SCI models have shown that anatomical brainstem changes can promote sensorimotor recovery.^40–43^ Furthermore, clinical interventions designed to re-engage top-down pathways, using combined action observation, motor imagery, and executed movement, can increase brainstem volume in individuals with chronic SCI.^44^ This highlights the potential for these corticocuneate projections to be targeted not only to support reafferentation but also to help mitigate secondary structural degeneration. Importantly, this potential may extend beyond spinal cord injury to other conditions involving disrupted afferent input, such as peripheral nerve injury or limb amputation.

Together, our findings show that the human brainstem and thalamic relay nuclei are not exclusively driven by peripheral input. Moreover, they demonstrate that top-down cortical modulation of early somatosensory relays can persist despite chronic sensory deafferentation and structural degeneration. These insights deepen our understanding of human somatosensory processing and highlight new avenues for rehabilitation targeting preserved top-down pathways.

## 4. Materials and Methods

### 4.1 Participants

We recruited 17 individuals with chronic cervical SCI (i.e., ≥6 months post-injury) and 21 age-, sex-, and handedness-matched able-bodied control participants. One individual with chronic cervical SCI was excluded after data collection, as their injury was confirmed to be thoracic rather than cervical, resulting in a final sample of 16 individuals with chronic cervical SCI (mean age ± s.e.m. = 52.4 ± 3.5 years; 1 female, 3 left-handers; see Table 1). A further control participant was excluded due to corruption of their fMRI data during acquisition, resulting in a final sample of 20 control participants (mean age ± s.e.m. = 50.8 ± 3.5 years; 2 females; 3 left-handers). There was no significant difference in age across groups (t_(34)_ = 0.33, p = 0.75).

**Table 1.** SCI participant demographic and clinical details.

| ID | Sex (M/F) | Age (years) | Time since injury (years) | AIS grade (A/B/C/D) | Neurological level of injury | GRASSP score (left/right hand) | Tissue bridges |
| --- | --- | --- | --- | --- | --- | --- | --- |
| PT01 | M | 35 | 7 | A | C4 | 0/3 | 0 |
| PT02 | M | 74 | 19 | D | C7 | 38/21 | - |
| PT03 | M | 60 | 36 | C | C5 | 26/24 | - |
| PT04 | M | 43 | 21 | A | C6 | 6/12 | 0 |
| PT05 | F | 70 | 7 | D | C5 | 32/32 | 3.0 |
| PT06 | M | 44 | 4 | D | C4 | 33/33 | 1.1 |
| PT07 | M | 70 | 29 | A | C4 | 7/14 | - |
| PT08 | M | 38 | 7 | A | C4 | 9/10 | 0 |
| PT09 | M | 55 | 13 | D | C4 | 25/23 | 3.1 |
| PT10 | M | 31 | 11 | C | C5 | 15/39 | 0.8 |
| PT11 | M | 59 | 37 | A | C7 | 30/33 | 0 |
| PT12 | M | 52 | 1 | D | C7 | 34/31 | 1.5 |
| PT13 | M | 59 | 1 | D | C1 | 40/33 | 3.4 |
| PT14 | M | 38 | 1 | D | C4 | 40/34 | 3.2 |
| PT15 | M | 69 | 9 | D | C2 | 34/33 | 3.3 |
| PT16 | M | 42 | 0.5 | D | C3 | 42/42 | 1.8 |

Sex: M = male, F = female; Age = age in years; Time since injury = Time since injury in years; AIS grade = American Spinal Injury Association (ASIA) impairment Scale grade defined based on the International Standards for Neurological Classification of Spinal Cord Injury (ISNCSCI): A = complete, B = sensory incomplete, C = motor incomplete, D = motor incomplete; Neurological level of injury = defined based on the ISNCSCI; GRASSP score = Graded Redefined Assessment of Strength, Sensibility and Prehension for the left and right hands; Tissue bridges = smallest width of summed ventral and dorsal tissue bridges on the midsagittal slice at the lesion level. Tissue bridge width could not be quantified for three participants due to metal artefacts and insufficient image quality (reported as “-”).

Individuals with chronic cervical SCI were eligible if they were 18-75 years old, had no MRI contraindications, were at least 6 months post cervical SCI, had no neurological or bodily impairments unrelated to SCI, and had the capacity to provide informed consent. Control participants were eligible if they had no MRI contraindications, no neurological disorders or hand impairments, and the capacity to provide informed consent. The control data included in this study were previously reported in Howell et al.^45^

Ethical approval was granted by the Kantonale Ethikkommission Zürich (EK-2018-00937), and all participants provided written informed consent prior to study enrolment. The study is registered at clinicaltrials.gov (NCT03772548).

### 4.2 Clinical characterisation

We used the International Standards for Neurological Classification of Spinal Cord Injury (ISNCSCI) to classify injury completeness and the level of neurological impairment. Upper-limb sensory and motor integrity was assessed with the Graded Redefined Assessment of Strength, Sensibility, and Prehension (GRASSP).^46^ Because GRASSP motor scores capture overall upper-limb function, including the shoulder and upper arm, we extracted hand-only sensorimotor sub-scores for analysis to better reflect task performance in the MRI scanner (see Table 1).

### 4.3 Experimental design

We tested whether overt or attempted hand movements after cervical SCI activate cortical and subcortical nodes along the somatosensory processing stream. Previous human fMRI studies have shown that attempting to move a paralysed, deafferented limb can still elicit activity in the sensorimotor system^15,47,48^, demonstrating that attempted movement offers a viable approach to probing central contributions to somatosensory processing in the absence of afferent drive.

We used a blocked fMRI design with three conditions: (1) left-hand open-and-closing, (2) right-hand open-and-closing, and (3) lip movement. Note that the lip movement condition was not analysed in the present manuscript, as this was part of another investigation into face somatopy following cervical SCI somatotopy (see Howell et al.^49^ for details). During scanning, participants fixated on a green cross, with task instructions (“left hand,” “right hand,” “lips,” and “rest”) displayed in white text. Each run comprised 12-second movement blocks followed by 6-second rest blocks, with each condition repeated eight times in a counterbalanced order. The fMRI protocol consisted of 4 runs, totalling 28 minutes and 48 seconds.

### 4.4 MRI data acquisition

#### Functional MRI

Functional MRI acquisition was performed according to our previously described protocol.^45^ In brief, data were acquired on a 3T Philips Ingenia system (Best, The Netherlands) using a 15- channel HeadSpine coil, except for six participants for whom a 32-channel coil was required due to space constraints. High-resolution T1-weighted images were collected to cover the brain and brainstem (sequence parameters: echo time (TE): 4.4ms; repetition time (TR): 9.3ms; flip angle: 8°; resolution: 0.75mm^3^), and task fMRI data were obtained with a partial-brain coverage EPI sequence, covering the brainstem, thalamus, and S1 (sequence parameters: singleshot gradient echo, TE: 26ms; TR: 2500ms; flip angle: 82°; reconstructed spatial resolution: 1.8 x 1.8 x 1.8 mm3; field of view (FOV): 210 x 187.6 x 70.2mm3; SENSE factor: 2.1; with 179 volumes per run).

To enable post-hoc physiological noise correction, we recorded respiratory motion using a chest bellow throughout the scans (sampling rate: 496 Hz). We also recorded oxygen saturation and cardiac pulse using a pulse oximeter attached to the second toe of each participant. All physiological devices were connected to a data-acquisition unit (Invivo; Philips Medical Systems, Orlando, Florida), which interfaced with a desktop computer running the corresponding recording software.

#### Quantitative MRI

In a separate session, qMRI data was acquired to assess macro- and microstructural markers of neurodegeneration. We used a previously established multiparameter mapping (MPM) protocol on a 3 Tesla Prisma MRI scanner (Siemens Healthineers, Erlangen, Germany) equipped with a 64-channel head/neck radiofrequency (RF) coil.^50–52^ The protocol comprised three 3D multi-echo fast low-angle shot (FLASH) gradient-echo sequences with different contrasts: proton density (PD), longitudinal relaxation time (T1), and saturation magnetisation transfer (MTsat) weighted MRI. We derived quantitative maps of the longitudinal relaxation rate (R1=1/T1), PD, MTsat, and the effective transverse relaxation rate (R2*=1/T2*). Images were reconstructed at an isotropic resolution of 1 mm³ (field of view: 224 × 256 mm², 176 partitions), with a total scan time of ∼ 18 minutes. Parallel imaging was applied in the anterior– posterior phase-encoding direction (GRAPPA 2 × 2) with partial Fourier 6/8. Readout bandwidth was 480 Hz/pixel. Acquisition parameters were as follows: for MT-weighted MRI, TR = 37 ms, flip angle 6°; for PD-weighted MRI, TR = 18 ms, flip angle 4°; and for T1- weighted MRI, TR = 18 ms, flip angle 25°. For all three modalities, six echo times (TE) were acquired (TE range between 2.46 and 14.76 ms). Additional RF reference scans were acquired for bias correction: B1⁺ maps using the actual flip-angle imaging method (TR = 2000 ms), and B1⁻ sensitivity maps from head and body coils (gradient-echo sequences, voxel size 4.0 × 4.0 × 4.0 mm³, TR = 3.8ms, flip angle 6°). Finally, to quantify spared tissue bridges, we acquired sagittal T2-weighted MRI of the cervical spinal cord at the lesion level (sequence parameters: TR = 3500 ms; TE = 84 ms; flip angle = 160°; echo train length = 14; spatial resolution = 0.3 × 0.3 × 2.5 mm; FOV = 220 × 220 × 55 mm.

### 4.5 MRI preprocessing and analysis

#### Functional MRI

Analysis of the MRI data was performed using FSL v6.0.4 (http://fsl.fmrib.ox.ac.uk/fsl), Matlab v9.12 (R2022a), FreeSurfer v6.0 (https://surfer.nmr.mgh.harvard.edu/), SPM12 (http://www.fil.ion.ucl.ac.uk/spm/), and Connectome Workbench v1.5 (https://www.humanconnectome.org/software/connectome-workbench).

To ensure consistent alignment of the hemisphere contralateral to the more impaired hand, we flipped the raw data along the x-axis for SCI participants whose GRASSP scores indicated greater impairment on the right side, as well as for their matched controls. Accordingly, comparisons between the left and right hands are defined throughout the manuscript in functional terms, i.e. the more and less impaired hand for SCI participants, and the non-dominant hand and dominant hand for controls.

Preprocessing was performed as previously described.^45^ We used FSL’s Expert Analysis Tool (FEAT) for brain extraction^53^, motion correction using FMRIB’s Linear Image Registration Tool (MCFLIRT)^54^, high-pass temporal filtering with a 90-second cutoff^55^, and spatial smoothing using a 2 mm full-width-at-half-max (FWHM) Gaussian Kernel. Physiological noise correction was performed using FSL’s Physiological Noise Modelling (PNM)^56^, which involved slice-wise assignment of cardiac and respiratory phases and the inclusion of 34 physiological regressors (4th-order cardiac and respiratory harmonics, 2nd-order interaction terms, and regressors for heart rate and respiration volume). Independent component analysis (ICA) was also implemented using the Multivariate Exploratory Linear Optimized Decomposition interface (MELODIC)^57^, with components classified as signal or noise based on their spatial maps, time series, and power spectra, following the guidelines of Griffanti et al.^58^

Image co-registration was performed in a series of sequential steps, with visual inspection after each transformation. Functional data were first aligned to each participant’s T1-weighted anatomical image using FMRIB’s Linear Image Registration Tool (FLIRT)^54^, with a mutual information cost function and 6 degrees of freedom, and subsequently refined with boundary-based registration (BBR)^60^. Each T1-weighted image was then normalised to the MNI152 1mm^3^ template using FMRIB’s nonlinear registration tool (FNIRT)^61^ with 12 degrees of freedom.

First-level general linear models (GLMs) were estimated in FSL FEAT using FMRIB’s Improved Linear Model (FILM)^62^ with local autocorrelation correction. Task regressors were convolved with FMRIB’s Linear Optimal Basis Sets (FLOBS)^63^, from which only the canonical hemodynamic response function (HRF) estimates were carried forward to higher-level analysis. Physiological noise and ICA-derived components were included as nuisance regressors. For each participant, we computed contrasts comparing each hand-movement condition to rest, as well as between-hand contrasts, i.e., dominant versus non-dominant hand for controls and less- versus more-impaired hand for cervical SCI participants. Individual runs were combined using a second-level fixed-effects analysis implemented with FMRIB’s Local Analysis of Fixed Effects.^62^ Given the small size of the brainstem nuclei of interest, runs with an absolute mean displacement exceeding 1mm were excluded to minimise motion-related confounds. In total, 7 of 144 runs were removed, with no more than one run excluded per participant.

To maximise anatomical alignment accuracy and intersubject registration, group-level statistical analyses for the cortex were performed on surface-based models^64^, and subcortical analyses of the brainstem and thalamus in volumetric MNI space. Univariate activation maps were generated using the second-level COPE maps for the between-hand contrasts, normalised to the 1mm3 MNI and fs_LR 32k brain templates. As spatial smoothing constraints in the brainstem violate assumptions underlying parametric Gaussian field-based inference^65^, all third-level analyses were performed using nonparametric permutation testing. Volumetric analyses were conducted using FSL’s RANDOMISE with threshold-free cluster enhancement (TFCE) and family-wise error (FWE) correction.^66,67^ One-sample t-tests were performed with the default parameters (5000 permutations; variance smoothing = 5 mm). Permutation testing was restricted to anatomical masks of the medulla and thalamus defined in MNI space using the Harvard–Oxford subcortical atlas.^68,69^ The brainstem mask was manually extended caudally to include the full extent of the cuneate nuclei and restricted to the medulla by removing axial slices superior to the inferior pontine sulcus.^70^ The probabilistic thalamus mask was thresholded at 50% to minimise ventricular overlap and binarised. Resulting TFCE p-maps were FWE corrected at p < 0.05.

Surface models were reconstructed from each participant’s T1-weighted image using FreeSurfer.^71–73^ The COPE maps from the second-level GLM were mapped onto each participant’s native cortical surface and then resampled to the fs_LR 32k standard mesh using Connectome Workbench.^74,75^ For each contrast, group-level statistical significance was assessed using FSL’s permutation analysis of linear models (PALM; i.e., a surface-based non-parametric data testing equivalent)^67^, with sign-flipping and 5000 permutations. The group z-statistic images were thresholded at z > 3.1, and FWE corrected at p < 0.01.

Finally, to quantify the extent to which overt and attempted hand-movement engaged the hand somatosensory nuclei, we extracted the mean z-value for left and right hand movement contrasted with rest within predefined anatomical regions of interest (ROIs)^45^, encompassing the ipsi- and contralateral cuneate nuclei, VPL nuclei, and S1 hand areas (see Figure 3A). For each region, ipsilateral values were computed as the average activity in the ROI ipsilateral to the moving hand, i.e., left hand-left ROI and right hand-right ROI, and contralateral values as the average activity in the ROI contralateral to the moving hand, i.e., left hand-right ROI and right hand-left ROI.

#### Quantitative MRI

The MPM data were preprocessed using the hMRI toolbox^76^ within the SPM12 framework (Wellcome Centre for Human Neuroimaging, UCL, London, UK; RRID: SCR_007037) in MATLAB (R2022a; MathWorks, Natick, MA, USA; RRID: SCR_001622), and visually inspected after each step. Images were first reoriented to each subject’s T1-weighted image using the auto-reorient function. We then estimated quantitative maps of R1, PD, MTsat, and R2* from the multi-echo acquisitions, with corrections applied for transmit-field (B1⁺) inhomogeneities using the acquired reference scans, and for receive-field (B1⁻), estimated from the ratio of head- and body-coil images^76,77^.

To assist with normalisation and voxel-based morphometry (VBM) analyses, we segmented the MTsat maps into grey matter (GM), white matter (WM), and cerebrospinal fluid (CSF) using enhanced tissue probability maps.^78^ A study-specific DARTEL template was then generated from tissue-class images, and deformation fields were applied to normalise the parametric maps to 1 mm^3^ MNI space.^79^ No spatial smoothing was applied to the quantitative maps to preserve anatomical precision in the small brainstem and thalamic nuclei.^21^

To visualise group-level macrostructural differences, we assessed voxel-wise changes across the whole brain grey- and white-matter tissue probability maps. Tissue class images were normalised to 1mm^3^ MNI space, modulated by their Jacobian determinants to preserve local volume^79^, and then smoothed with a 2 mm full-width at half maximum (FWHM) Gaussian kernel. The VBM analyses were then performed in SPM12 using an independent two-sample t-test design, with age, sex, and total intracranial volume (TIV; GM + WM + CSF) included as covariates of no interest.^80^ Clusters were first defined using a voxelwise threshold of *p* < 0.001 (uncorrected). Statistical significance of these clusters was then determined using cluster-level family-wise error (FWE) correction at *p* < 0.05.

To quantify tissue microstructure, we used MTsat and R2*. MTsat is sensitive to tissue lipid and protein content and is considered an in vivo marker of myelin integrity^81–84^, whereas R2* is sensitive to iron content and broader microstructural properties of tissue organisation^85,86^. To characterise these properties and their relationship to fMRI activity, we extracted mean MTsat and R2* values from the same anatomically defined ROIs used in the univariate fMRI analysis. For each region, values were averaged across the left and right homologous ROIs.

#### Midsagittal tissue bridge analysis

Lesion severity at the spinal injury site was quantified using midsagittal tissue bridge measurements that have been previously reported for this SCI cohort.^49^ Tissue bridges were defined as hypointense intramedullary regions separating the hyperintense cerebrospinal fluid from the cystic cavity on midsagittal slices. The smallest ventral and dorsal bridge widths were measured and summed to yield the total tissue bridge width. Images from three participants were excluded due to metal artefacts or insufficient image quality that precluded reliable quantification.

### 4.6 Statistical analysis

Statistical analyses were conducted in R (v4.4.0; RStudio) and visualised using Graphpad Prism (v.10.5.0). Linear mixed-effects models were implemented using the *lme4*^87^ and *lmerTest*^88^ packages. Model assumptions were verified by visual inspection and simulation-based diagnostics using *DHARMa.*^89^ Importantly, residuals were approximately normally distributed and homoscedastic, with no evidence of overdispersion or influential outliers for the final analysis. Additionally, all analyses were repeated without outlier values exceeding ±3 SD (i.e., one observation in the control group for each of the cuneate and S1 ROI analyses) to confirm that they did not alter any statistical conclusions.

For the ROI analyses, fixed effects included Group (SCI, control) and Side (ipsilateral, contralateral), with Participant entered as a random intercept. Significant main or interaction effects were followed by two-tailed post hoc pairwise contrasts (*emmeans*) with false discovery rate (FDR) correction. Effect sizes were reported as partial η² for omnibus effects and Cohen’s d for independent group comparisons, computed using the *effectsize* package^91^. Additionally, we tested planned contrasts between the Group and Side differences, correcting for multiple comparisons within each ROI. Brainstem activation was tested against zero using one-sample t-tests (FDR corrected), and case-control comparisons were made using Crawford–Howell modified t-tests^92^. For quantitative MRI analyses (MTsat and R2*), group differences were tested separately within each ROI using Welch’s two-sample t-tests.

Given the small sample size (n = 16), we used Spearman rank correlations to test the structure– function relationships within each ROI. All analyses were two-tailed with α = 0.05. In addition, clinical variables were examined as predictors of structural and functional measures using multiple regression models. Multicollinearity was assessed using variance inflation factors, with all predictors showing acceptable levels. Due to their exploratory nature, no correction for multiple comparisons was applied here.

Bayesian analyses were performed using the *BayesFactor* package.^93^ Bayes factors (BF10) were interpreted according to Dienes^94^: BF_10_ < 1/3 indicates substantial evidence for the null hypothesis; BF_10_ > 3 indicates substantial evidence for the alternative hypothesis; and values between 1/3 and 3 indicate weak or anecdotal evidence. Bayes factors were computed using Bayesian model comparisons for main and interaction effects and Bayesian t-tests for paired or independent post hoc contrasts.

## Data Availability

Data analysis was performed using publicly available toolboxes and script templates, with modifications as described in the Methods section of the manuscript. Researchers wishing to replicate or build on this work are encouraged to contact the corresponding author for further guidance. The raw data analysed in this study cannot be shared due to confidentiality restrictions outlined in the ethics agreement approved by the Ethics Committee of the Canton of Zürich (KEK-2018-00937). However, aggregated and anonymised group-level data are available upon request to support reproducibility.

## Acknowledgements

We thank the subjects for their participation in the study. We are grateful to Sijamini Baskaralingam, Adrian Taubner and Charlotte Meninghin for their help with data collection, and to Simon Schading-Sassenhausen for his support with the tissue bridge dataset. We thank Roger Luchinger, the Institute for Biomedical Imaging and the Swiss Center for Musculoskeletal Imaging for scanning support.

