## Supplementary Material for "Top-down processing alone activates the early somatosensory nuclei"

**Supplementary Methods**

**EMG analysis**

Most SCI participants in our study had incomplete injuries and retained some hand-movement ability, meaning they still received residual bottom-up afferent input during the fMRI task. In contrast, one participant (PT01) was clinically complete, with total paralysis of both hands and only minimal residual cutaneous sensation over the right thumb. This case subject provided an opportunity to verify that activation in our regions of interest can be elicited in the absence of any afferent input. To confirm that no peripheral muscle activity was elicited in PT01 during attempted hand movements, we recorded surface electromyography (EMG) in a separate session outside the MRI scanner. The same procedure was carried out in a matched control participant. Under the premise that PT01 is unable to generate overt movement, we can exclude confounding bottom-up afferent input to supralesional somatosensory regions. This would indicate that any observed brain activation during the fMRI would instead reflect top-down processes.

Participants performed the same task as in the fMRI session. Bilateral surface EMG recordings (Trigno Wireless, Delsys) targeted the primary muscles driving finger flexion and extension: extensor digitorum (ED), extensor pollicis (EP), flexor digitorum (FD), the first dorsal interosseous (FDI), and the thenar eminence (TE) (Supplementary Figure 1). Since voluntary contractions could not be used to verify electrode placement, we positioned the electrodes using anatomical landmarks and fibre orientation, following established surface EMG guidelines (SENIAM: http://seniam.org/arm_location.htm; Axelgaard Anatomy Atlas: https://www.axelgaard.com/App/Anatomy/Finger).

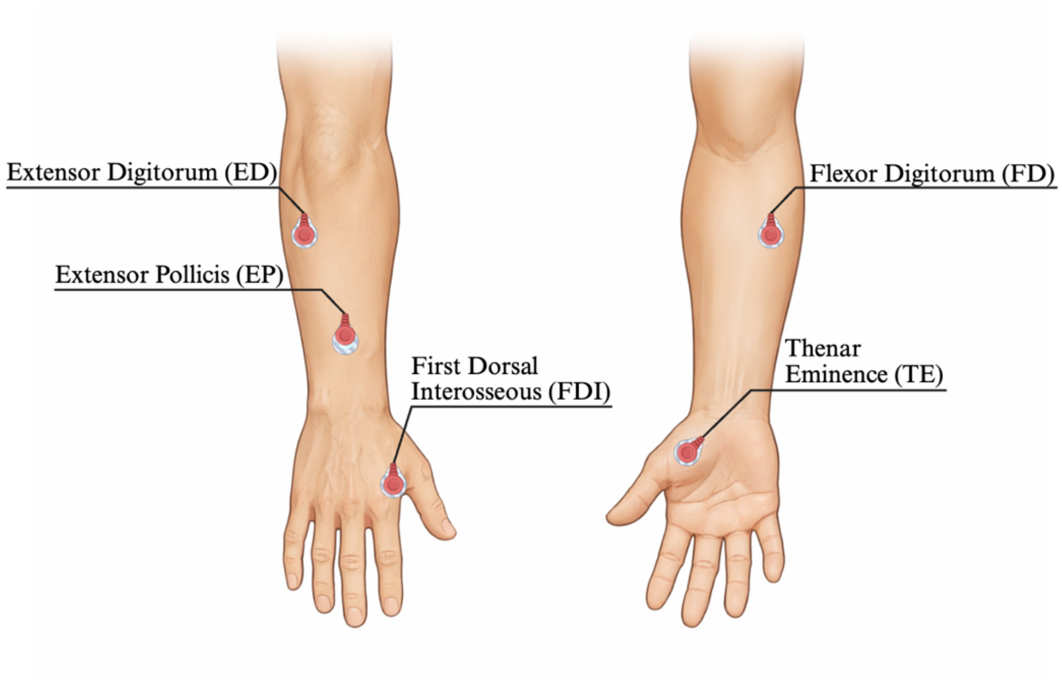


**Supplementary Figure 1. Surface electromyography electrode placement**

We recorded surface electromyography (EMG) during overt (HC01) and attempted movement (PT01). Electrodes were placed bilaterally over the key muscles responsible for finger flexion and extension: extensor digitorum (ED), extensor pollicis (EP), first dorsal interosseous (FDI), flexor digitorum (FD), and the thenar eminence (TE). Because voluntary contractions could not be used to verify sensor location, placement was guided by anatomical landmarks and fibre orientation following established surface EMG guidelines (SENIAM; Axelgaard Anatomy Atlas).

EMG data preprocessing for each muscle included offset correction using a 100-ms baseline interval extracted during the initial rest period, high-pass filtering at 30 Hz, and notch filtering at 50 Hz. The EMG signal traces were then divided into the 16 movement and 16 rest blocks, excluding the first second of each period to account for delays in movement initiation. Finally, the root-mean-square (RMS) amplitude was calculated for each segment. As only one control and one SCI participant were recorded, statistical comparisons were performed at the single-subject level. Given the non-normal distribution and unequal variances of the data, we used Wilcoxon signed-rank tests to compare trial-wise EMG activity during rest and movement per muscle. We used MATLAB's filloutliers to check for and replace outliers in the RMS values. In total, four trials were identified (two from HC02 and two from PT01). P-values were corrected across muscles using false discovery rate correction for multiple comparisons.

**No peripheral motor output during attempted movement in complete SCI**

To characterise the expected level of peripheral motor output during the task, we first analysed EMG recordings from the control participant. All sampled muscles, except the right flexor digitorum, showed a clear increase in EMG amplitude during overt finger movements compared with rest, confirming that the task reliably elicits peripheral motor activity in an intact system (Supplementary Figure 2A; Right ED: $W = 136$, $p < 0.01$; Right EP: $W = 136$, $p < 0.01$; Right FD: $W = -24$, $p = 0.11$; Right FDI: $W = 136$, $p < 0.01$; Right TE: $W = 136$, $p < 0.01$; Left ED: $W = 136$, $p < 0.01$; Left EP: $W = 136$, $p < 0.01$; All Left $|W| = 136$, $p < 0.01$, p-values FDR corrected). The absent flexor digitorum response in the right arm is likely attributable to both the muscles' deep location and reliance on anatomical landmarks rather than muscle contraction for the electrode placement. In contrast, none of the recorded muscles in the paralysed limbs of PT01 showed any increase in EMG amplitude during attempted movement (Supplementary Figure 2B; Right ED: $W = 24$, $p = 0.95$; Right EP: $W = 2$, $p = 0.99$; Right FD: $W = -24$, $p = 0.95$; Right FDI: $W = 58$, $p = 0.73$; Right TE: $W = -12$, $p = 0.99$; Left ED: $W = -44$, $p = 0.35$; Left EP: $W = 62$, $p = 0.35$; Left FD: $W = -44$, $p = 0.35$; Left FDI: $W = 56$, $p = 0.35$; Left TE: $W = 32$, $p = 0.44$, p-values FDR corrected). This absence of peripheral motor output indicates that the participant generated no overt muscle activity capable of producing proprioceptive or mechanoreceptor-driven afferent signals. Accordingly, any task-evoked responses observed in PT01's fMRI data must reflect top-down cortical drive rather than peripheral input (Supplementary Figure 2C).

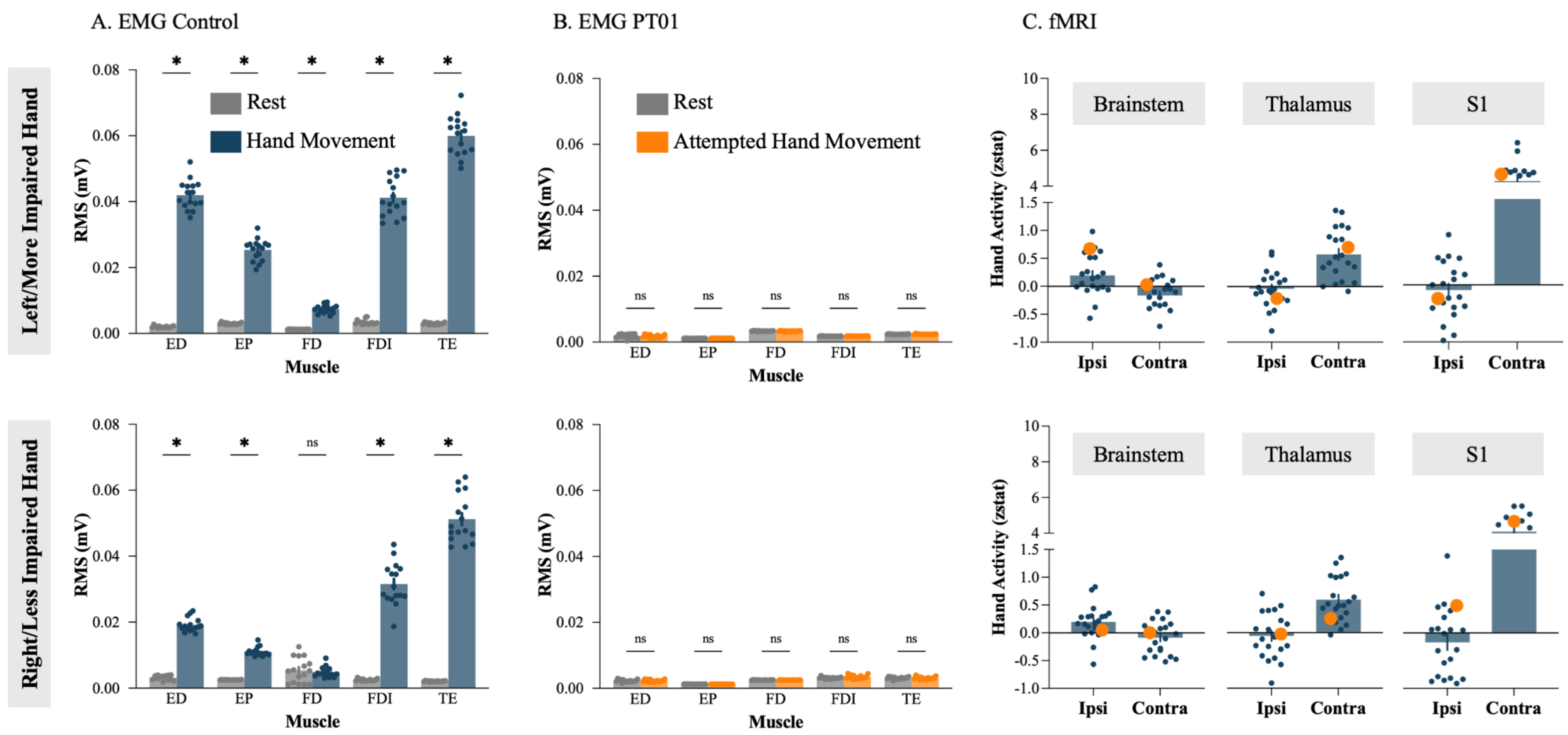

A. EMG Control
B. EMG PT01
C. fMRI
Left/More Impaired Hand
Right/Less Impaired Hand
Rest
Hand Movement
Attempted Hand Movement
RMS (mV)
Muscle
ED
EP
FD
FDI
TE
ns
*
Hand Activity (zstat)
Brainstem
Thalamus
S1
Ipsi
Contra

**Supplementary Figure 2. No increase in peripheral motor output during attempted hand movement in complete SCI**

To determine whether fMRI responses in PT01 could arise from bottom-up afferent signals, we examined whether the hand-movement/attempted-movement task elicited peripheral motor output in a control participant and a clinically complete SCI participant (PT01). In the control participant (A), significant increases in EMG amplitude were observed during hand movement (blue) relative to rest (grey) across all recorded muscles, except the right flexor digitorum (FD), confirming that the task reliably generates peripheral motor output in an intact system. In contrast, none of the recorded muscles in the clinically complete SCI participant (B; PT01) exhibited any increase in EMG amplitude during attempted hand movement (orange) compared with rest (grey), demonstrating the absence of overt muscle activity and associated afferent signals. Accordingly, the preserved laterality (C) observed in PT01's fMRI responses (orange dots) along the somatosensory pathway must reflect top-down cortical drive rather than residual peripheral input. The control fMRI responses (blue dots) reflect both bottom-up and top-down somatosensory signal processing. ED = extensor digitorum; EP = extensor pollicis; FD = flexor digitorum; FDI = first dorsal interosseous; TE = thenar eminence; RMS = root mean square; mV = millivolt. Error bars represent ± standard error of the mean; p-values are false discovery rate-corrected for multiple comparisons. * = $p \leq 0.01$, ns: non-significant.

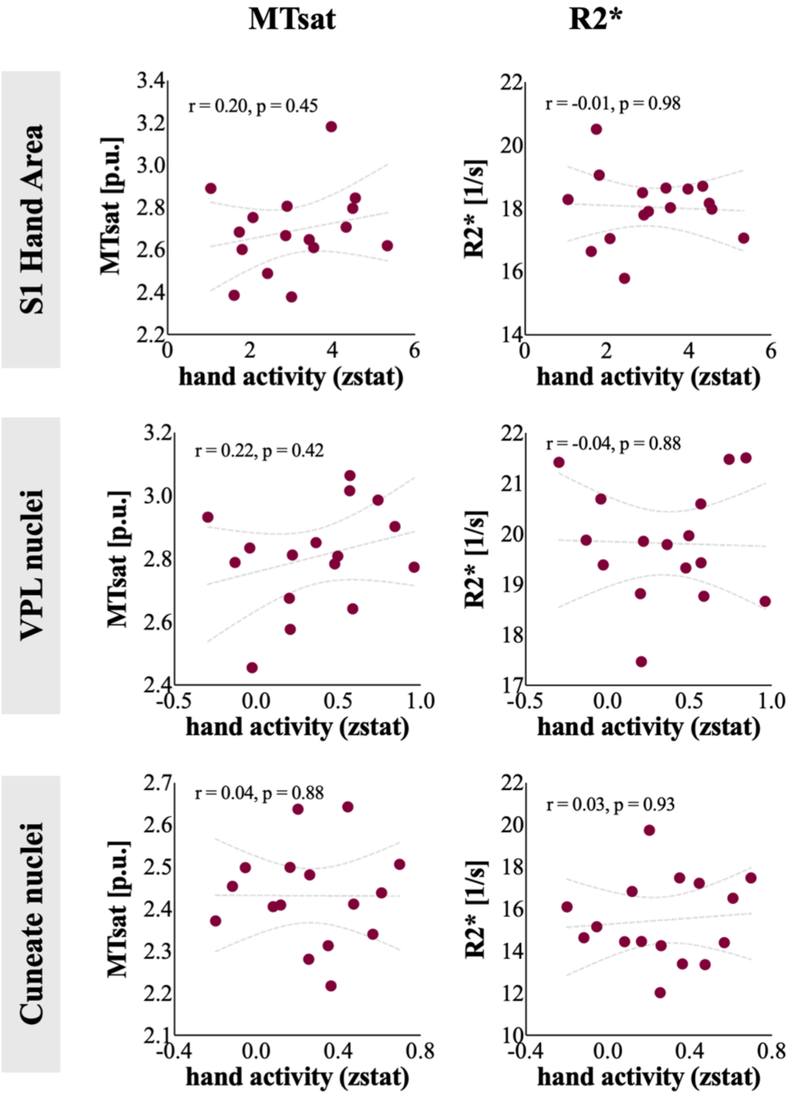


**Supplementary Figure 3: Relationship between task-evoked activation and microstructural integrity across somatosensory relay nuclei**

We examined whether the magnitude of task-evoked activity during hand movement/attempted movement was associated with local tissue microstructure along the somatosensory pathway in cervical SCI participants. Spearman correlations were computed between the mean activation estimates (zstat) in the ipsilateral cuneate nucleus, the contralateral ventroposterior lateral (VPL) thalamus, and the contralateral S1 hand area, and quantitative MRI measures of microstructural tissue integrity (MTsat and R2*). Across all regions and metrics, no significant associations were observed, suggesting that preserved task-evoked activity was not linearly related to the degree of microstructural degeneration. Individual SCI participants are plotted as red circles; dashed lines indicate best-fit regression lines with 95% confidence bands. MTsat values are expressed in percentage units (p.u.), and R2* values in per-second units (1/s). P-values shown are uncorrected.

**A. Full Models**

| ROI | Outcome | $R^2$ | Predictor | B | SE B | p-value |
|---|---|---|---|---|---|---|
| *Cuneate* | **MTsat** | 0.56 | GRASSP | 0.004 | 0.001 | **0.021*** |
| | | | Time since injury | 0.005 | 0.003 | 0.121 |
| | | | Tissue bridge | -0.033 | 0.028 | 0.276 |
| | **R2*** | 0.55 | GRASSP | 0.04 | 0.025 | 0.142 |
| | | | Time since injury | 0.107 | 0.052 | **0.07#** |
| | | | Tissue bridge | -0.737 | 0.477 | 0.156 |
| | **fMRI activity** | 0.19 | GRASSP | 0.001 | 0.004 | 0.693 |
| | | | Time since injury | 0 | 0.007 | 0.949 |
| | | | Tissue bridge | 0.048 | 0.068 | 0.498 |
| *VPL* | **MTsat** | 0.12 | GRASSP | 0.002 | 0.002 | 0.467 |
| | | | Time since injury | -0.002 | 0.005 | 0.622 |
| | | | Tissue bridge | -0.047 | 0.043 | 0.308 |
| | **R2*** | 0.31 | GRASSP | 0.024 | 0.017 | 0.207 |
| | | | Time since injury | -0.023 | 0.036 | 0.542 |
| | | | Tissue bridge | -0.672 | 0.334 | 0.075 |
| | **fMRI activity** | 0.04 | GRASSP | 0.002 | 0.006 | 0.71 |
| | | | Time since injury | -0.006 | 0.013 | 0.673 |
| | | | Tissue bridge | -0.023 | 0.12 | 0.853 |
| *S1* | **MTsat** | 0.29 | GRASSP | 0.003 | 0.003 | 0.287 |
| | | | Time since injury | -0.009 | 0.005 | 0.142 |
| | | | Tissue bridge | -0.066 | 0.049 | 0.218 |
| | **R2*** | 0.17 | GRASSP | -0.012 | 0.018 | 0.528 |
| | | | Time since injury | 0.001 | 0.038 | 0.983 |
| | | | Tissue bridge | -0.153 | 0.347 | 0.67 |
| | **fMRI activity** | 0.06 | GRASSP | -0.002 | 0.023 | 0.922 |
| | | | Time since injury | -0.005 | 0.047 | 0.916 |
| | | | Tissue bridge | 0.236 | 0.432 | 0.597 |

**B. Reduced Models**

| ROI | Outcome | $R^2$ | Predictor | B | SE B | p-value |
|---|---|---|---|---|---|---|
| *Cuneate* | **MTsat** | 0.47 | GRASSP | 0.003 | 0.001 | **0.023*** |
| | | | Time since injury | 0.006 | 0.002 | **0.013*** |
| | **R2*** | 0.33 | GRASSP | 0.01 | 0.02 | 0.628 |
| | | | Time since injury | 0.097 | 0.039 | **0.027*** |
| | **fMRI activity** | 0.02 | GRASSP | 0.002 | 0.003 | 0.643 |
| | | | Time since injury | 0 | 0.006 | 0.995 |
| *VPL* | **MTsat** | 0.10 | GRASSP | 0.001 | 0.002 | 0.621 |
| | | | Time since injury | -0.003 | 0.004 | 0.409 |
| | **R2*** | 0.01 | GRASSP | 0.003 | 0.014 | 0.833 |
| | | | Time since injury | 0.007 | 0.027 | 0.794 |
| | **fMRI activity** | 0.14 | GRASSP | 0.002 | 0.004 | 0.592 |

| | | | | | | |
|---|---|---|---|---|---|---|
| | | | Time since injury | -0.009 | 0.008 | 0.283 |
| ***S1*** | **MTsat** | 0.40 | GRASSP | 0.001 | 0.002 | 0.587 |
| | | | Time since injury | -0.009 | 0.004 | **0.023*** |
| | **R2*** | 0.12 | GRASSP | -0.016 | 0.013 | 0.217 |
| | | | Time since injury | -0.015 | 0.024 | 0.544 |
| | **fMRI activity** | 0.07 | GRASSP | 0.003 | 0.015 | 0.832 |
| | | | Time since injury | -0.025 | 0.028 | 0.398 |

**Supplementary Table 1. Multiple regression analyses of clinical predictors across somatosensory regions of interest.** Multiple regression analyses testing whether clinical measures predict structural (MTsat, R2*) and functional (mean zstat) indices across somatosensory regions of interest (cuneate nucleus, VPL, and S1). Full models (A) included retained hand function (GRASSP), injury chronicity (time since injury), and tissue bridge width (full models), with reduced models (B) excluding tissue bridge width due to missing data. $R^2$ values reflect the variance explained by the model for each ROI and outcome. Unstandardised coefficients (B), standard errors (SE·B), and p-values are reported. *p < 0.05; #p < 0.10.
